# Environmental change drives multi-generational shifts in the gut microbiome that mirror changing animal fitness

**DOI:** 10.1101/2023.10.24.563854

**Authors:** Candace L. Williams, Claire E. Williams, Shauna N. D. King, Debra M. Shier

**Author notes:** Contributed equally (Candace L. Williams, Claire E. Williams). Corresponding author: Candace L. Williams, Conservation Science, San Diego Zoo Wildlife Alliance, 15600 San Pasqual Valley Road, Escondido, CA, USA.

## Abstract

Gut microbiomes can dramatically affect host health and fitness and can also be highly dynamic in response to changing environmental conditions. This intricate interplay between gut microbiota, environmental pressures, and host health necessitates accounting for all these variables when predicting the response of animals to a changing environment. These predictions are of broad concern but are highly relevant to conservation biology, and more specifically to populations transitioning from the wild to human care for the purposes of *ex situ* breeding programs. Captivity can dramatically alter both host-associated microbial communities and host health, but the dynamics of how the host and microbiome transition between the wild and captive state—within and across generations—are not well described. Here, we evaluate the microbiome and host fitness metrics for Pacific pocket mice (*Perognathus longimembris pacificus*) during the establishment of an *ex situ* conservation breeding program to characterize how pocket mice and their microbiomes respond to the environmental pressures of captivity across five generations. We found that the microbiome assumes a novel, stable conformation after two to three generations in captivity and that the patterns observed in the transitioning microbiota mirror the patterns of transitioning hosts’ weight and reproductive metrics. Moreover, we identify several microbial taxa which are correlated with successful reproduction. Our results provide insight not only into the effects of captivity but can be broadly applicable for understanding the effects of environmental change on organisms and their gut microbiota, which we propose may require multiple generations to reach an alternative stable state.

## Introduction

Human induced rapid environmental change negatively affects native species through habitat loss, fragmentation, and degradation (1); see reviews by (2–5). Animals that live within these ecosystems must disperse, acclimate, or adapt to persist, but for many species, their options are limited. The degree of habitat loss and/or fragmentation may prevent successful dispersal, and acclimatization may be difficult depending on the type and magnitude of environmental perturbations (*e.g*., invasive species, pollutants, climate change) or inherent characteristics of the animal affected (*e.g*., dispersal capacity, genetic diversity). Given the rapid nature of these environmental changes, many animals cannot evolve quickly enough to adapt and, therefore, human intervention is required for their persistence. One strategy used by wildlife managers to prevent imminent extinction is the creation of conservation programs that couple captive breeding and habitat restoration (6, 7), where animals are brought into human care, propagated, and offspring are released into suitable wild habitats. However, selective pressures differ between wild and captive environments (*e.g.*, altered environment and diet) (8, 9), and captive adaptations may be maladaptive when animals are reintroduced into the wild (10).

Microbiome-mediated plasticity may assist animals in acclimatizing or adapting to environmental change (11, 12). Since microbiomes are highly dynamic, they can change on a more rapid timescale than their animal hosts in response to environmental pressures (12–19). Gut microbiota, in particular, play many vital roles in host physiology (20–23), and environmental change may lead to an imbalance in the microbiome, which can have negative outcomes for hosts. Specifically, captivity has been shown to alter the gut microbiome, as observed by the reduction in microbial community richness and shifts in composition (24), as well as increased associations with detrimental effects on host health including gastrointestinal disease and metabolic disorders (25–32). It is the cumulative effect of these captivity-associated perturbations on both the host and its microbial community which can ultimately shape the health and survival of animals with the potential to impact the success of breeding programs. However, the dynamics of how the host and microbiome transition between the wild and captive environments are not well understood. In order to make effective management decisions, it may be vital to understand how the gut microbiome mediates host fitness (33). Detailed characterization of these dynamics could enable interventions targeted at the gut microbiome (*e.g*., dietary intervention and/or transfaunation, see (34–36)) and lead to improved success of conservation breeding programs, reintroductions, and ultimately the recovery of endangered species.

The intricate interplay between gut microbial communities, environmental pressures, and host health necessitates the inclusion of all these variables when predicting the response of animals to a changing environment. Yet, most studies have been cross-sectional in design, sampling only established populations in captivity and those in the wild or monitoring a transition over a short time scale (37). Here, we evaluated how environmental change altered the gut microbiome and associated animal fitness in the endangered Pacific pocket mouse (*Perognathus longimembris pacificus*) across multiple generations. The Pacific pocket mouse is a species in the heteromyid family that was previously thought to be extinct in the 1970s but rediscovered in 1993 and emergency listed as endangered (38). Due to the precarious status of the three remaining extant populations, a conservation breeding program was initiated in 2012 whose goal was to create three new populations through reintroduction (39).

Using 16S rRNA amplicon sequencing in combination with detailed monitoring of the health and reproductive status of pocket mice, we observed the dynamics of simultaneous host and microbial response to *ex situ* conditions of founding individuals as they transitioned to captivity and across descendant matrilineal generations maintained in captivity. By combining both host and bacterial measures, we aimed to better understand the bidirectional interaction between the microbiome and pocket mice undergoing rapid environmental change. We tested the hypothesis that the gut microbiome would change as animals adapted to the *ex situ* environment, and that these changes would persist through multiple generations. Further, we predicted that changes in the gut microbiome elicited by the transition to captivity would correspond to changes in pocket mouse health and fitness across multiple generations.

## Methods

All analyses and associated sample metadata described below can be found in our GitHub repository (https://github.com/clw224/2023-Williams-PPM-WildtoCaptive/) as well as a visual description of our workflow in Figure S1.

### Sample collection

We collected fecal samples from Pacific pocket mice (*n =* 129 individuals) either from trap, enclosure, or scruff across multiple matrilineal lineages (F_0_-F_4_), with some repeat measurements for wild-collected (F_0_) individuals, from 2012-2020 (totaling *n =* 184 samples). When trapping extant sites *in situ*, we inspected traps between animals to ensure the fecal pellets removed from the trap during processing belonged to the target animal. Using sterilized tweezers, we transferred feces into a vial. For *ex situ* collection, we used sterilized tweezers to remove feces from different areas of the enclosure and transferred them to a vial. We also collected some samples directly from pocket mice *in* and *ex situ*, as they often defecate when being handled; therefore, when handling the pocket mice by scruff, we collected the feces directly into a vial. We stored samples at -20 °C prior to analysis.

### Animal fitness data collection

To evaluate fitness metrics *in* and *ex situ*, we collected data on weights and reproductive readiness. In 2007 and 2008 we conducted trapping *in situ* five nights each week after pocket mice emerged from hibernation (February or March), through the active season (∼September) each year. We marked pocket mice for individual identification using Visible Implant Elastomer (Northwest Marine Technology, Inc. Shaw Island, WA, USA) injected in unique color combinations just under skin on the sides of the tail base. During processing, we weighed each individual and assessed reproductive status. For females we evaluated reproductive status by visually evaluating vulvar swelling. We rated the degree of vulvar swelling on a four point scale (1 = not swollen, 4 = maximally swollen) developed for kangaroo rats (40) and adapted for pocket mice. For males we categorized reproductive status using testes position (*i.e.,* 0 = non-scrotal, 1 = partially scrotal, and 2 = scrotal). We used the same methods to collect fitness metrics *ex situ* except that for pocket mice housed in the *ex situ* population, we obtained weekly weights for all pocket mice throughout the year. We assessed reproductive status for all females at least every 3 days during the active season. If females had a swelling of ≥ 2, we monitored them as often as every hour to determine physical estrus (41). We used a swelling score of ≥ 3, perforation, or presence of a plug to indicate that an estrous cycle had occurred. We assessed male reproductive condition weekly. To evaluate parentage, we tracked pocket mouse pedigrees through the studbook in PopLink (42) and used PMx to manage breedings (43).

### DNA extraction

We lyophilized fecal samples (*n =* 184) and homogenized ∼3 mg of fecal pellets in 80 % methanol in water to extract metabolites for a separate, unpublished dataset (both, Optima LC-MS grade, Fisher Chemical). Following metabolite extraction, we used the remaining cell pellet for genomic DNA extraction. In addition to samples, each 96-well plate contained both negative and positive controls for DNA extraction and PCR (ZymoBIOMICS Microbial Community Standard and ZymoBIOMICS Microbial Community DNA Standard, respectively, Zymo Research, Irvine, CA, USA), with samples and controls randomized across both extraction plates. We mechanically lysed samples by two bead-beating steps (5 min), with a heat treatment (65 °C, 10 min) in between, and lysate was further extracted using a Zymo Fecal, Soil, Microbe 96 Magbead kit (Zymo Research, Irvine, CA, USA) on an OT-2 liquid handling robot (Opentrons, Queens, NY, USA) using a protocol modified from Sanders *et al.* 2022. We quantified the resulting genomic DNA by Quant-it Assay (Invitrogen, Waltham, MA, USA).

### Sequencing library preparation

We prepared sequencing libraries using a protocol modified from Kozich *et al.* (2013) and Williams *et al.* (2019). Briefly, we PCR amplified the V4 region of the 16S rRNA gene in a 25-μl reaction mixture with 1× KAPA HiFi Hot Start Ready mix (Kapa Biosystems, Wilmington, MA, USA), 0.2 μM was added of each primer, and sample concentrations ranged between 0.5 to 10 ng genomic DNA. Amplification conditions were as follows: 95 °C for 2 min, followed by 25 cycles of 95 °C for 20 s, 55 °C for 15 s, and 72 °C for 30 s and a final 10-min extension at 72 °C. We purified PCR products via gel extraction (Zymo Gel DNA recovery kit; Zymo Research, Irvine, CA, USA) using a 1.0 % low-melt agarose gel (National Diagnostics, Atlanta, GA, USA). While the negative controls produced no band, we excised the expected area. We quantified purified PCR products by Quant-it assay (Invitrogen, Waltham, MA, USA). We combined all samples to yield an equimolar 2 nM pool. Following the manufacturer’s protocol, we sequenced using an Illumina MiSeq reagent kit v2 (500 cycles, (Illumina, San Diego, CA, USA) with the final library concentration of 3.5 pM with 20 % Phi-X spiked in.

### 16S rRNA sequence analyses

We used QIIME2 and R as described previously by Williams, Brown and Williams (2023) for sequence analysis. In brief, we imported demultiplexed FASTQ files into QIIME2 (47), and denoised (trimmed and filtered) using the DADA2 plug-in, with command “denoise-paired” to generate ASVs (amplicon sequence variants) (48). We built a phylogenetic tree by alignment using the MAFFT plug-in, masking using the FastTree plug-in, and rooting it via the phylogeny midpoint plug-in (49, 50). We assigned taxonomy using the feature-classifier “classify-sklearn” with the SILVA full-length 16S database (reference database release 138) (51). Although the negative controls had few reads (< 100), we decontaminated the ASV feature table in R via the *decontam* package (52), and we filtered any ASVs which were not gut-associated from the feature table in QIIME2. We also removed sequences assigned to mitochondria, chloroplasts, or those that lacked phylum-level classifications to avoid issues associated with misamplification.

### ASV analyses

To quantify compositional beta diversity and determine if it varied across generations, we performed a robust Aitchison Principal Component Analysis (RPCA) on the ASV table implemented by the DEICODE plugin in QIIME2 (53). We found that when tracking changes across generations, significant differences in beta diversity variance emerged (*P =* 0.002, via PERMDISP). To accurately quantify differences in bacterial composition, we randomly trimmed our dataset to provide a balance of sample observations across generations (See Table S1) based on several criteria: 1) F_0_ populations, samples were grouped by F_0_-wild (samples taken at animal collection *in situ*), F_0_-captive (following the 3-6 month transition period *ex situ*), 2) if duplicate samples existed for individuals, they were grouped to create one sample per individual, 3) we randomly selected individuals across generations, but took into account sex, to ensure even sampling across males and females (F_0_, *n =* 19 [F_0_-wild, *n =* 11; F_0_-captive *n =* 16]; F_1_, *n =* 17; F_2_, *n =* 29; F_3_, *n =* 31; F_4_, *n =* 24).

Our statistical analyses of microbiome data fell into two comparisons: 1) determining the changes within a wild-caught individual as they adapted to captivity (F_0_ only), and 2) evaluating the change in the gut microbiome across generations of mice, from F_0_ through F_4_. To evaluate the transition of bacterial alpha diversity as pocket mice adapted to captivity, we calculated Shannon diversity using QIIME2 and constructed a linear mixed effects model with time as a fixed effect and individual mouse identification as a random effect (54–56). To test for differences in Shannon diversity across generations we used a simple linear regression. To understand how beta diversity changed from the wild through the transition to captivity and across generations, we used the DEICODE distance matrix and measured differences between groups and their dispersion (variance) using PERMANOVA (permutational analysis of variance; Anderson 2017) and betadisper implemented by *vegan::adonis and vegan::betadisper,* respectively, in R (58), and for calculations regarding adaptation to captivity, we used individual pocket mouse ID as a random effect. We also examined the relationship between the first principal coordinate (PC1) and time in captivity using a Pearson correlation (*stats::cor*) (59).

To determine which ASVs or bacterial families comprised a stable, “core” community that was maintained across generations, we used an abundance-occupancy approach (60). Briefly, ASVs and their families were ranked based on their occupancy and abundance in each generation. We used a less than 2.0% contribution to Bray-Curtis community composition as a cut off as suggested by Shade and Stopnisek (60) and Bray-Curtis similarity in our dataset plateaued shortly after this cut-off (61). Therefore, we likely included the majority of ASVs that are important to structuring the community beta diversity, thus identifying the taxa which are maintained over generations and are therefore considered potentially important for the community structure of the gut microbiome. To identify which taxa were differentially abundant across generations, we used *Maaslin2* (62) in R to implement linear models to assess differences in relative abundance of taxa across generations with a Benjamini-Hochberg multiple comparison correction (63), as it provides a good balance between discovery of statistically significant ASVs and limitation of false positive occurrences.

### Host fitness analyses

To test for effects of generations in captivity on pocket mouse fitness, we used weight, estrous cycles, or testes score as a response variable and generation as a fixed effect, and compared changes in these reproductive measures by generation using Kruskal-Wallis tests due to lack of normality (59, 64). For all outcome variables, we calculated a mean for each individual across their first full breeding season (either the year following their capture or their birth in captivity). For estrous cycles we censored pocket mice for which other variables may confound cycling (*e.g.*, pregnancy). To quantify the number of offspring produced by each pocket mouse, the package *purgeR* was used on pedigrees generated from the studbook. To test for relationships between specific microbial taxa on pocket mice reproductive success, we constructed separate generalized linear models implemented with the package in *Maaslin2* (62). We rarefied our ASV table to 8500 reads in QIIME2 (47), and then we used pocket mouse reproduction (*i.e*., pups born) in their first breeding season (0/1) as a binomial response variable and relative microbial abundance as fixed effect, correcting for multiple comparisons using a Benjamini-Hochberg approach (63).

### Data availability

Upon publication of this manuscript, raw sequences and corresponding metadata will be available in the NCBI Sequencing Read Archive. All other relevant data, protocols, and scripts, including all statistical models used for analysis, will also be made available in a public GitHub repository: https://github.com/clw224/2023-Williams-PPM-WildtoCaptive/.

## Results

### Captivity rapidly alters microbiome composition

To understand how the gut microbiome changes during the transition to and throughout subsequent generations in captivity, we evaluated the community composition and richness of the gut microbiome across these periods (Figure 1a). We found that the pocket mouse gut microbiome’s community composition (beta diversity) changed rapidly as wild-collected individuals transitioned to captivity (PERMANOVA, *P =* 0.004; Figure 1b). The influence of transitioning to captivity on microbial community composition is particularly apparent along the first principal component (PC1; Figure 1b). By contrast, richness and evenness (alpha diversity) remained relatively stable across the first year in captivity, with no significant differences observed across time periods (*P =* 0.13; Figure 1d). By comparing the composition of the pocket mice gut microbiome across generations in captivity, we observed how dramatic this shift is for wild-born animals relative to subsequent captive-born generations, with significant shifts again observed along PC1 (Figure 1c, Figure S2a). However, significant compositional changes in subsequent captive-born generations are observed along PC2, with each generation being significantly different from the previous, excluding F_3_-F_4_ (PERMANOVA, *P =* 0.0012, *P =* 0.017, *P =* 0.0012, *P =* 0.10; respectively; Figure S2b). Interestingly, we observed the opposite relationship with dispersion (PERMDISP; *P =* 0.033). Across F_0_-wild to F_3_ we see no significant difference in compositional variance across these generations (PERMDISP, all *P >* 0.38), but we observed differences in variance between both wild-born pocket mice and F_2_ generations compared to F_4_ (PERMDISP, both *P =* 0.015). Taken together, compositional shifts that are occurring across F_0_-F_3_ in combination with similarly high levels of dispersion across generations prior to F_4_, suggests that the microbiome community composition stabilized after F_3_ (Figure 1c, S1b). Despite the observed shifts in the pocket mouse gut microbiome composition across generations, there was no corresponding change in the level of alpha diversity (*P =* 0.09) (Figure 1e).

**Figure 1.**
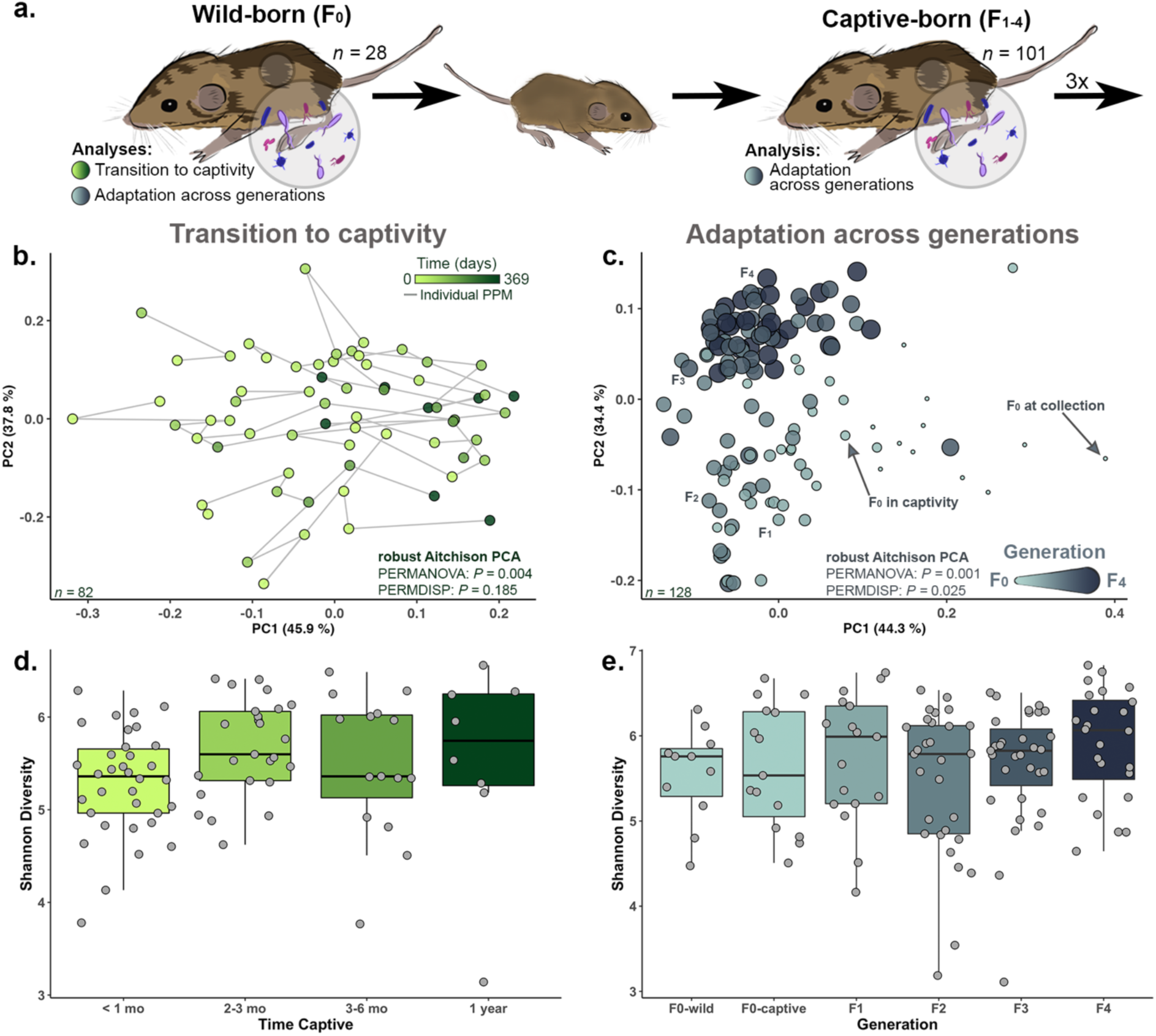
Captivity significantly alters microbiome composition across multiple generations, but not alpha diversity. (a) An overview of our experimental design, sampling from both wild-born (F_0_) and captive-born (F_1-4_) individuals in the conservation breeding program. A robust Aitchison principal component analysis (b) displayed that microbiome composition shifted as F_0_ individuals transitioned to captivity, while Shannon diversity (d) remained relatively constant. Community composition showed further changes across generations (c) while Shannon diversity did not (e).

### Microbial extirpation and replacement occur through several generations in captivity

Across all pocket mice sampled, we found the most dominant bacterial phyla were Firmicutes (mean relative abundance = 62.2 %), Bacteroidota (29.3 %), Cyanobacteria (3.10 %), Proteobacteria (2.10 %), Actinobacteriota (1.70 %), and Verrucomicrobiota (1.50 %). At the family level, the dominant six taxa were classified as Lachnospiraceae (24.5 %), Muribaculaceae (21.7 %), Erysipelotrichaceae (15.3 %), Rikenellaceae (7.00 %), Ruminococcaceae (5.70 %), and Clostridia - UCG-014 (3.60). Most ASVs which were shown to be differentially abundant across generations were members of microbial families which were present across all generations based on abundance-occupancy analyses. The proportion of the gut microbiota contributed by these core families remained relatively constant across generations (Figure 2a), while ASVs within those families underwent an extirpation and replacement process (Figure 2b-e). Some ASVs which dominated the community in wild animals were almost completely lost in F_1+_ individuals. For example, a Lachnospiraceae ASV and a Streptococcaceae ASV were both replaced by novel ASVs within the same bacterial family (Figure 2c, Figure 2e, respectively). In other cases, one dominant ASV in wild pocket mice, a member of the Muribaculaceae, was replaced by several lower abundance taxa (Figure 2b), while multiple ASVs within the Clostridia family change across generations (Figure 2d).

**Figure 2.**
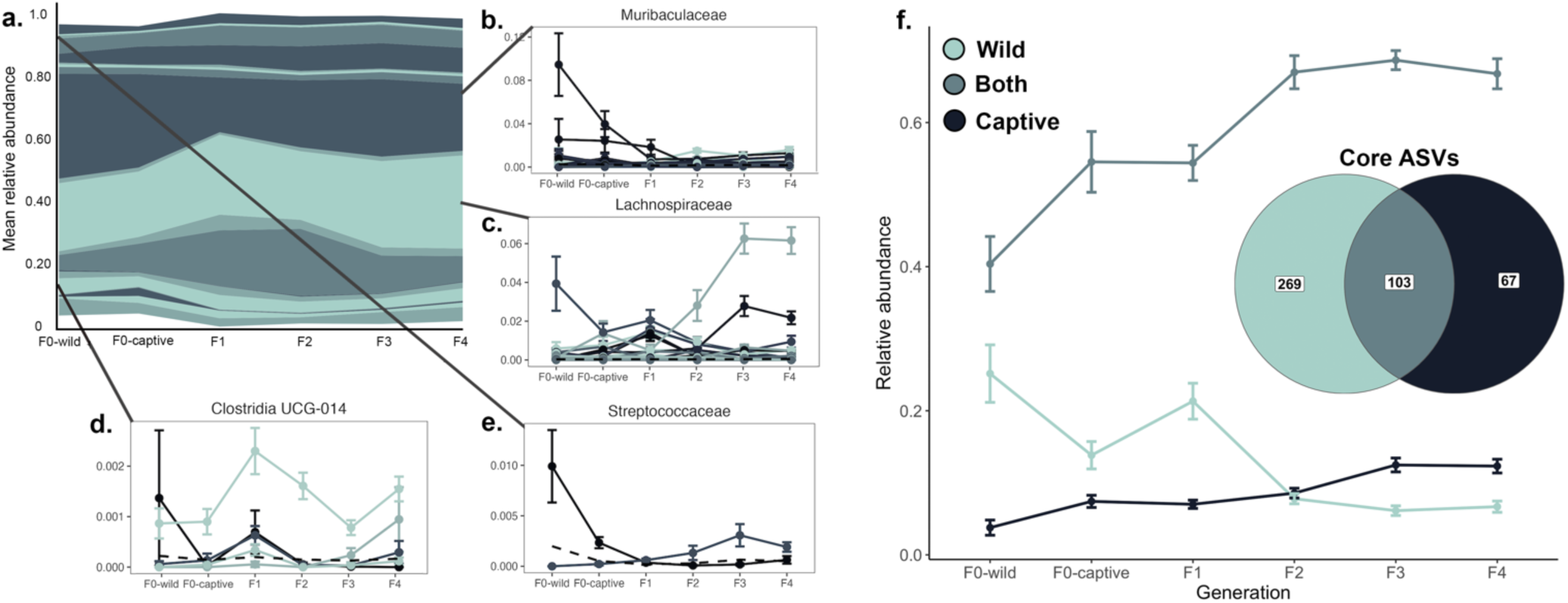
The mean relative abundance of core microbial families remained relatively constant across generations while some individual ASVs within those families underwent extirpation and replacement. (a) Stream graph displaying the mean relative abundance of each family deemed core by an abundance-occupancy analysis across generations. The width of each bar illustrates the mean proportion of the core microbiome composed of that family in each generation. (b-e) display ASVs which exhibited significant differences in mean relative abundance across generations, illustrating that while families remained constant, individual ASVs showed substantial flux. (f) The pocket mice core microbiome shifts to a new conformation - including different ASVs after several generations in captivity (inset Venn diagram). The ASVs considered core in wild animals (light blue) decrease in mean relative abundance and are replaced by ASVs which are core in captive animals (dark blue). Error bars represent standard error.

While the core families in the PPM gut microbiota remained constant, the ASVs considered “core” to a wild mouse versus “captive” mouse (F_3+_) only partially overlapped (Figure 2f). In total, 372 ASVs were considered core to the wild population at collection, with ASVs only associated with wild-core grouping comprising ∼ 25 % of the total microbial community, compared to 170 ASVs, with captive-core grouping ASVs comprising ∼15 % of the F_3+_ population. However, 103 taxa that were shared by both core groups were observed at 40 - 65 %. Many ASVs which contributed to the core microbiota of wild pocket mice were lost, as the relative abundance of ASVs considered part of both the wild and captive core increased, with cross-over between the two core-groupings occurring in the F_2_ generation (Figure 2f). Despite these changes, we found that the total relative abundance of the core communities for each grouping to be relatively stable (F_0_-wild core proportion: 69.3 %; F_4_ core proportion: 85.7 %). This shift to a novel core community corresponds to the stabilization observed in the gut microbial community composition.

### Animal fitness measures mirror microbiome dynamics

Conservation breeding programs rely on healthy, reproductively fit animals to produce offspring destined for reintroduction, and managers often use mean weight and reproductive status as proxies for evaluating wildlife health. Here, we examined these measures across the entire *ex situ* population within the conservation breeding program from its inception (F_0_-F_6_) and used data collected *in situ* from extant populations as a baseline for comparison. We observed a transition in mean weight across generations in captivity (Figure 3a and 3c, Kruskal Wallis *Χ^2^* =32.515 (Males), 28.493 (Females); *P <* 0.001 for males and females) where both male and female F_0_ in captivity have a significantly higher weight which declines and approaches the wild baseline after F_3_ and beyond (Wilcoxon *P <* 0.01 for F_0_ vs F_3_ comparison in both males and females). This approach to wild baseline is also observed in the reproductive metrics of pocket mice transitioning to captivity (Figure 3b and 3d, Kruskal-Wallis *Χ^2^* = 16.489 (Males) and 12.304 (Females); *P =* 0.01 (Males) and *P =* 0.056 (Females)). F_0_ and F_1_ females have fewer estrous cycles per month, and estrous cycling increases toward the wild baseline, stabilizing in F_4_ individuals and beyond. Although these differences are not statistically significant after correcting for multiple comparisons, this stabilization pattern mirrors what we observed with the microbiome data. Both pocket mice fitness and microbiome measures exhibit a clear period of flux through F_2_, after which point these metrics stabilize (*i.e*., pocket mice adopt a novel, but stable microbial community in F_3_ and beyond, while reproductive metrics stabilize close to the wild baseline). Moreover, the relative abundance of some ASVs predicted pocket mouse reproductive fitness (Figure 4, all q-values for reproduction < 0.05). Despite the substantial flux we observed in the gut microbiome, these taxa did not change significantly from generation F_0_ through generation F_4_ (all q-values for generational differences > 0.05).

**Figure 3.**
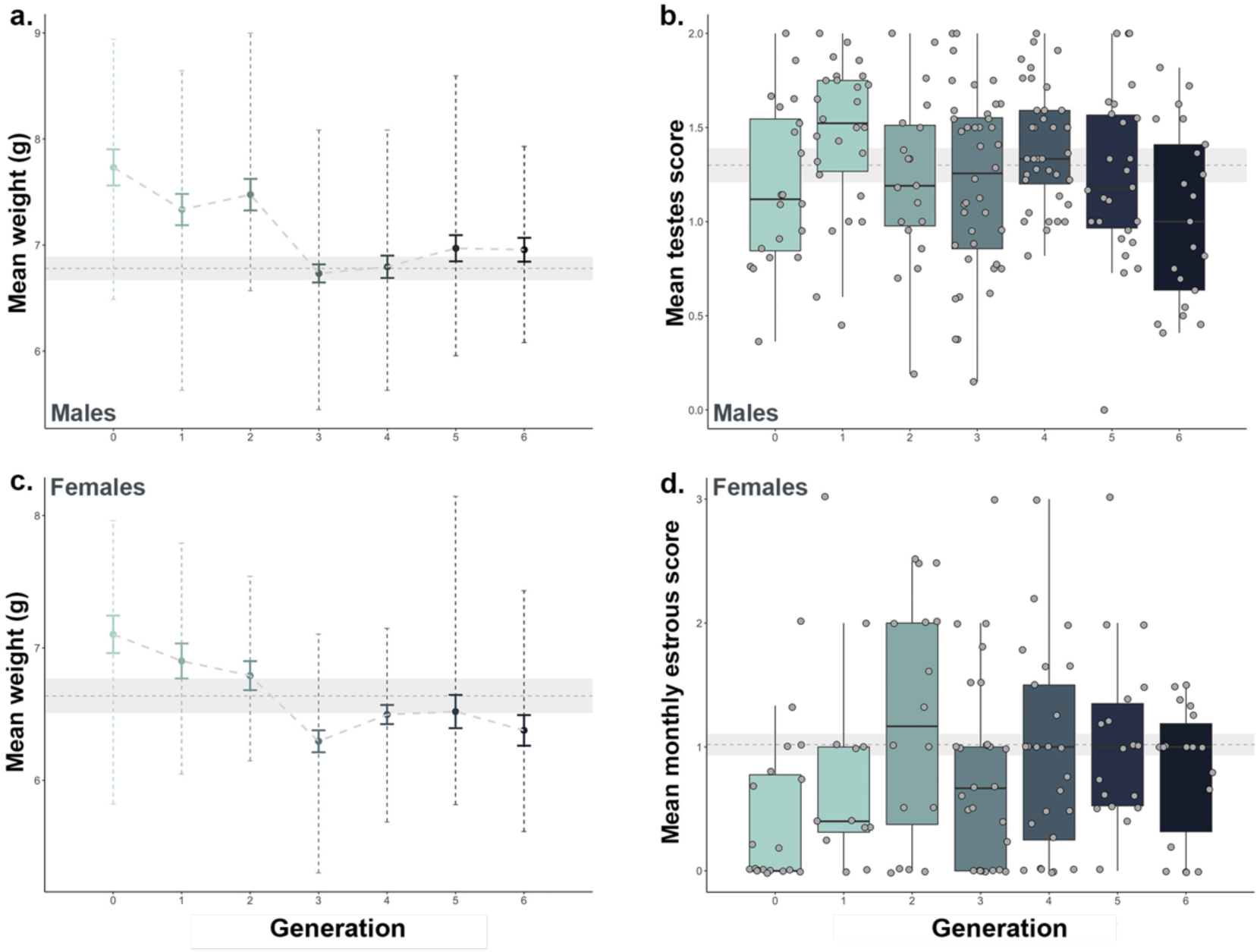
Fitness metrics change across pocket mice generations. In the first generation, weights of both males (a) and females (c) and male testes score (b) and female monthly estrous score (d) compared to wild baseline (shown in gray, mean ± standard error baseline levels). Error bars represent standard error.

**Figure 4.**
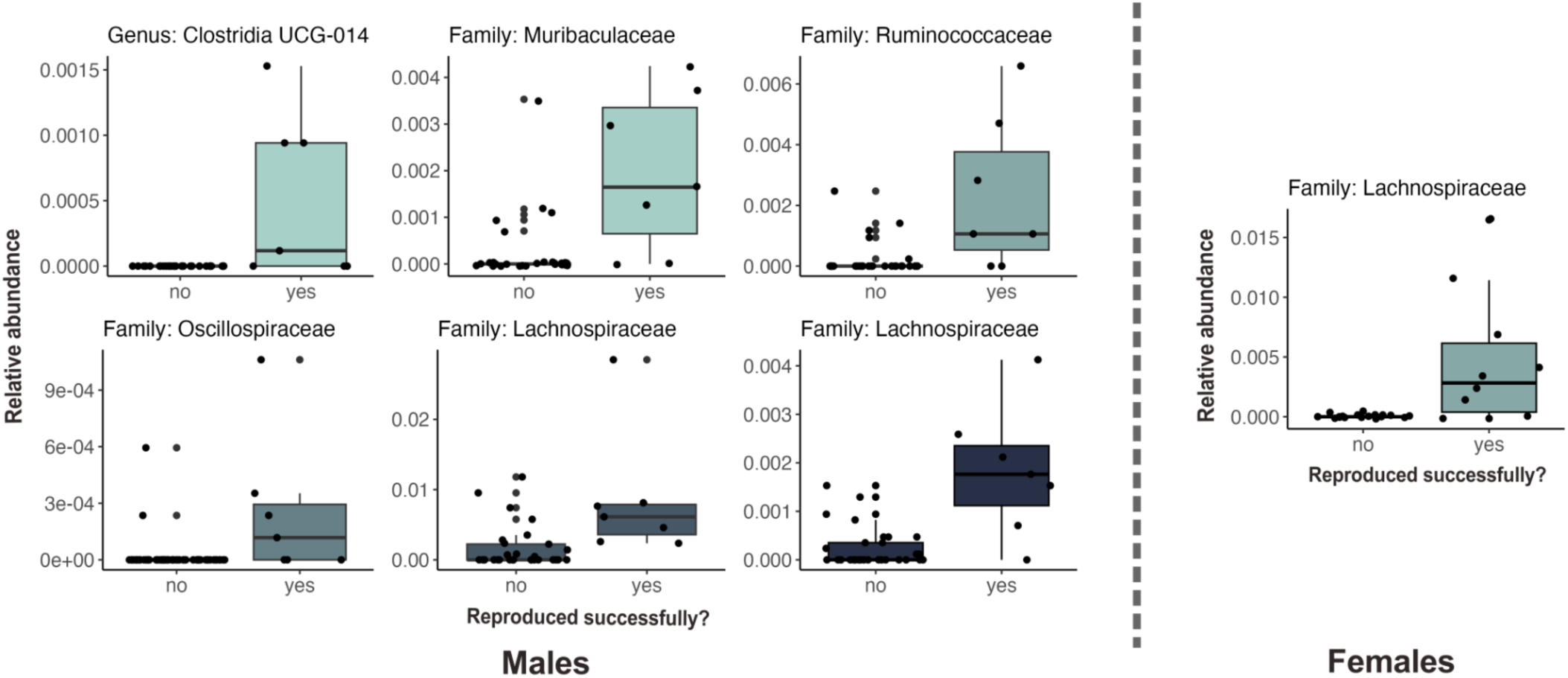
Relationship between microbial taxa and successful reproduction in the first full breeding season. Taxa associations are divided by males (left) and females (right). Each boxplot displays the relative abundance of each associated ASV, with the lowest taxonomy annotation available indicated in the plot title.

## Discussion

As pocket mice transitioned to captivity, their gut microbiome community composition shifted significantly, both within the first generation, and through generation F_2_, after which no further significant changes occurred, indicating that a “captive” microbiome may have emerged in F_3+_ animals. Yet, these significant changes to microbiome community composition occurred despite no significant changes in alpha diversity. Reduction in alpha diversity is often noted as a hallmark of captivity when compared to wild populations (24, 65), yet most comparative studies lack sampling longitudinally to observe how individuals transition between *in* and *ex situ,* which may bias the results. With our repeat sampling, we observed a reduction in dispersion in the gut microbiota, indicating that while the individual-level alpha diversity of the microbiota does not change, the population-level range of variation is reduced in captive pocket mice. This pattern has also been observed in other studies indicating that reduction in diversity may indeed be characteristic of captivity (35, 66, 67), though whether it manifests at the individual or population level may vary among species (24, 68). Moreover, pocket mice in our *ex situ* population receive native seed enrichment and live in a facility with skylights that allow for natural photoperiod and enclosures that allow for the retention of many of their natural behaviors, such as burrowing in sandy-soil substrates and living solitarily but with visual and olfactory conspecific cues. It is possible that one reason we may not see a significant loss of richness is through this more successful mimicking of their natural habitats (69, 70). Regardless, as alpha diversity remains constant, shifts in community composition occur due to changes in the relative abundance of taxa or the even extirpation and replacement of individual microbial ASVs.

Our study revealed that the mean relative abundances of core microbial families in pocket mice stayed relatively constant across generations, indicating that these families may serve important health and fitness functions for the species. These included taxa that are typical members of the mouse (*Mus* spp.) microbiome, including Muribaculaceae, which has been linked to complex carbohydrate utilization (71–73); Lachnospiraceae, a gut symbiont involved in nutrient acquisition and energy production (74, 75); Streptococcaceae, which are linked to memory and learning (72, 76); and Clostridia which are associated with host metabolic homeostasis (77). However, we observed a significant change in the relative abundance of many different ASVs within those core microbial families. This observation indicated that extirpation and replacement of microbes within core families is likely a major driving force behind the compositional changes observed in the pocket mice gut microbiota, but due to a potentially high functional redundancy within some bacterial families, may not necessarily result in significant fitness tradeoffs (78, 79).

To understand if these extirpation and replacement dynamics led to a novel captive core community, we compared the ASV-level core microbiota of wild pocket mice to that of pocket mice in generations F_3_ and F_4_ (after the microbial community had stabilized, referred to as the “captive” microbiome). While many core ASVs overlapped, we observed a loss of many wild-core ASVs alongside an increase in captive-core ASVs, indicating that there is a portion of the gut microbiota which may be more substantially influenced by the environment, leading to population-level differentiation. It is unclear what the exact source of these novel ASVs is. While diet is certainly a driver of the changes in the gut microbiome in captivity, interaction with humans, the built environment, or a combination of these factors may also contribute novel taxa (68). Altogether, the shift between a wild and captive core microbiota but not a change in their proportion in the overall microbiome, coupled with high intra-generational compositional variance during the transition between wild and captive microbiota, may indicate a dynamic change between alternative stable states in the gut microbiome (80).

We documented a substantial deviation from the wild baseline values for both weight and estrous cycling in early generations in captivity. However, by generation F_2_-F_3_ we observed a transition back to the wild baseline for these metrics followed by a stabilization. This occurs along the same timeline as the stabilization observed in the gut microbiota. Not only do we find similar stabilization trends between microbiome and host fitness metrics, but we also found the presence of several microbes to be predictive of successful male and female reproduction in their first year, including three Lachnospiraceae ASVs. Several studies have shown links between the gut microbiome and reproductive physiology, including microbial transformation of endogenous hormones or their endocrine-disrupting chemical mimics, such as phytoestrogens (31, 81–83). More specifically, members of Lachnospiraceae have been associated with increased fertility in captive female southern white rhinoceros (Williams *et al.,* 2019) and reproductive hormone cycling, including a negative correlation to fecal progesterone metabolites in captive female black rhinoceros (84). As progesterone concentrations are typically low in early follicular and pre-ovulatory phases while estrogen is high (85) an inverse relationship may also be inferred. In other words, a negative correlation of Lachnospiraceae ASVs with progesterone may equate to a positive correlation to pre-ovulatory phases prior to estrus.

Taken together, the simultaneous changes in gut microbiota and reproductive measures may provide further evidence of a bidirectional connection between the gut microbiome and reproductive physiology. Specifically, transition to captivity may induce changes in the gut microbial community, leading to an unstable transition state which might negatively affect host physiology (80). After several generations in captivity, a novel, but healthy, alternative stable microbiome state may emerge and support host reproductive health and physiology. If so, then the gut microbiome may be able to rapidly respond to environmental pressures and assume a healthy stable state which is suitable for the present host environment (15, 16, 86, 87). Simply put, fitness effects conferred by some microbial taxa may be context dependent.

Our results illustrate how pocket mice and their microbial symbionts adapt to environmental change and provide insight into how the species may also respond when transitioning from *ex* to *in situ* upon reintroduction, indicating that close monitoring of populations through the first few generations could be important given implications that changes in the gut microbiome due to the environmental changes induced by translocation may alter fitness. Alternatively, negative effects on host physiological processes mediated by the stress of captivity may lead to an altered gut microbiome. While several studies have linked the gut microbiome to endocrine function (reviewed in 22), it remains a possibility that the gut microbiome is simply an indicator of physiological stress, which responds, alongside many other aspects of host physiology, to rapid environmental change. Additional studies examining changes in the gut microbiome and reproductive physiology could help elucidate the causative nature of this relationship.

While our results provide significant insight into the dynamics of the gut microbiome as it responds to captivity within one and across multiple generations, several of our results should be interpreted with caution. First, due to the small population sizes of wild pocket mice, our conservation breeding program is composed of a small number of host genotypes (88), and reproduction was low across several early generations. Therefore, it remains difficult to disentangle the effects of host genetic variation versus environment on the gut microbiome and reproductive metrics in our population. This is an area of active debate (*e.g*., 88, 89), and future work could attempt to parse out the effects of genetic variation and the gut microbiome on host response to environmental change using data collected in conservation breeding programs. Regardless, it is likely that *both* host genetic variation and gut microbiome composition play a role in host health and fitness and may need to be considered together for future conservation management decisions (91).

As anthropogenic activity continues to alter ecosystems across the globe and threaten the persistence of many species, human interventions like conservation breeding programs are becoming increasingly important to protect biodiversity. We know that the gut microbiome can respond to the pressures of captivity and also impact host fitness (29, 31, 92, 93). Thus, understanding how the gut microbiome may impact the success of these programs - both upon entering captivity and upon reintroduction – may influence recovery of at-risk species. More broadly, our results elucidate the effects of environmental change on organisms and their gut microbiota. We illustrate that environmental perturbations can result in an altered, highly variable gut microbiome which persists for multiple generations. However, eventually, the gut microbiome can stabilize alongside host metrics in the novel environment, providing much hope for the long-term success of such conservation programs. How transitions between stable states of the gut microbiome, and their tipping points, might be directly linked with host fitness remain open questions. Regardless, gut microbiota appear to be an important piece of the puzzle when understanding how animals may respond to rapid changes in their environment.

## Supplemental information

**Figure S1.**
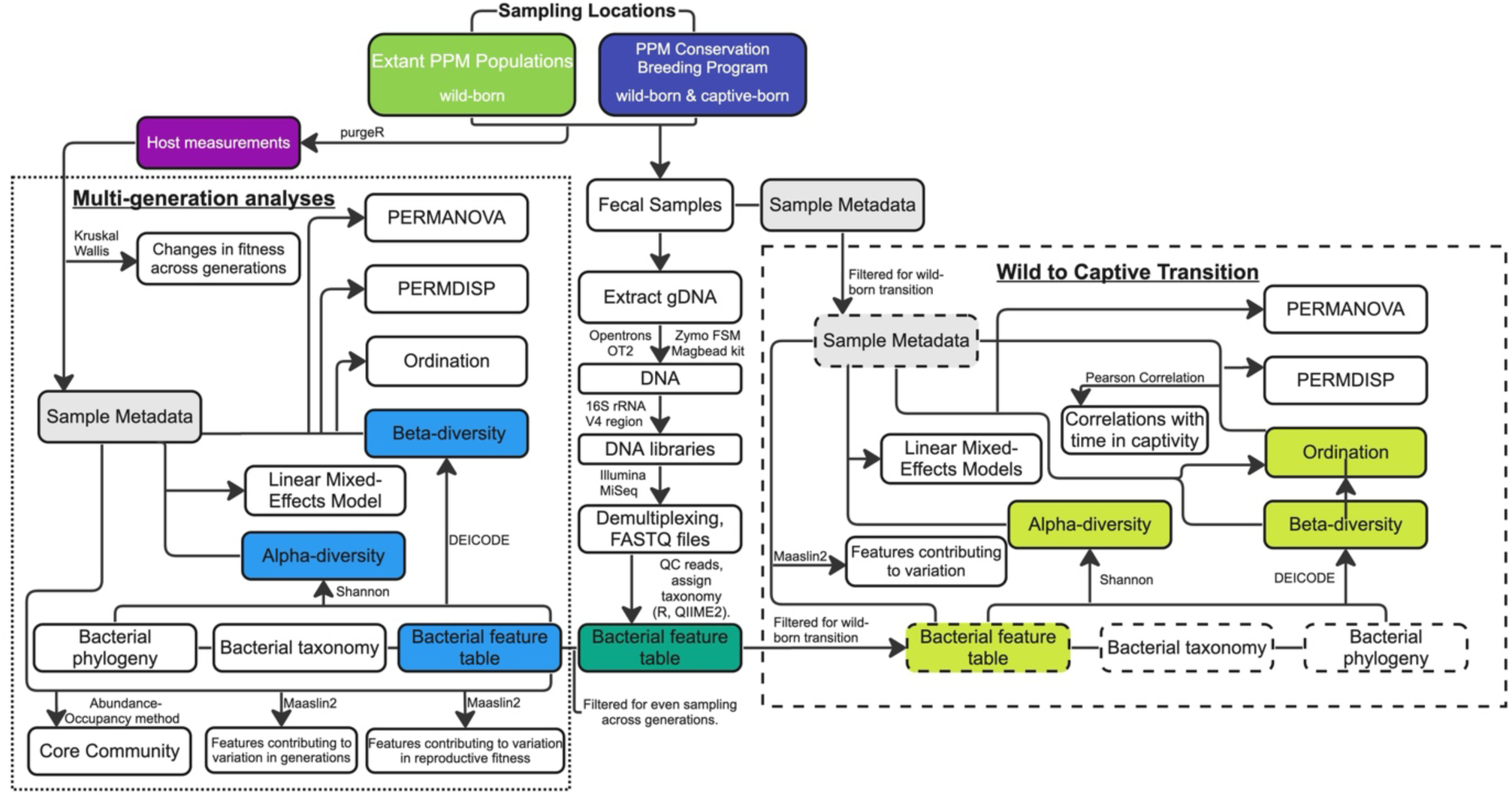
Visual workflow of all methods and analyses conducted for this manuscript.

**Figure S2.**
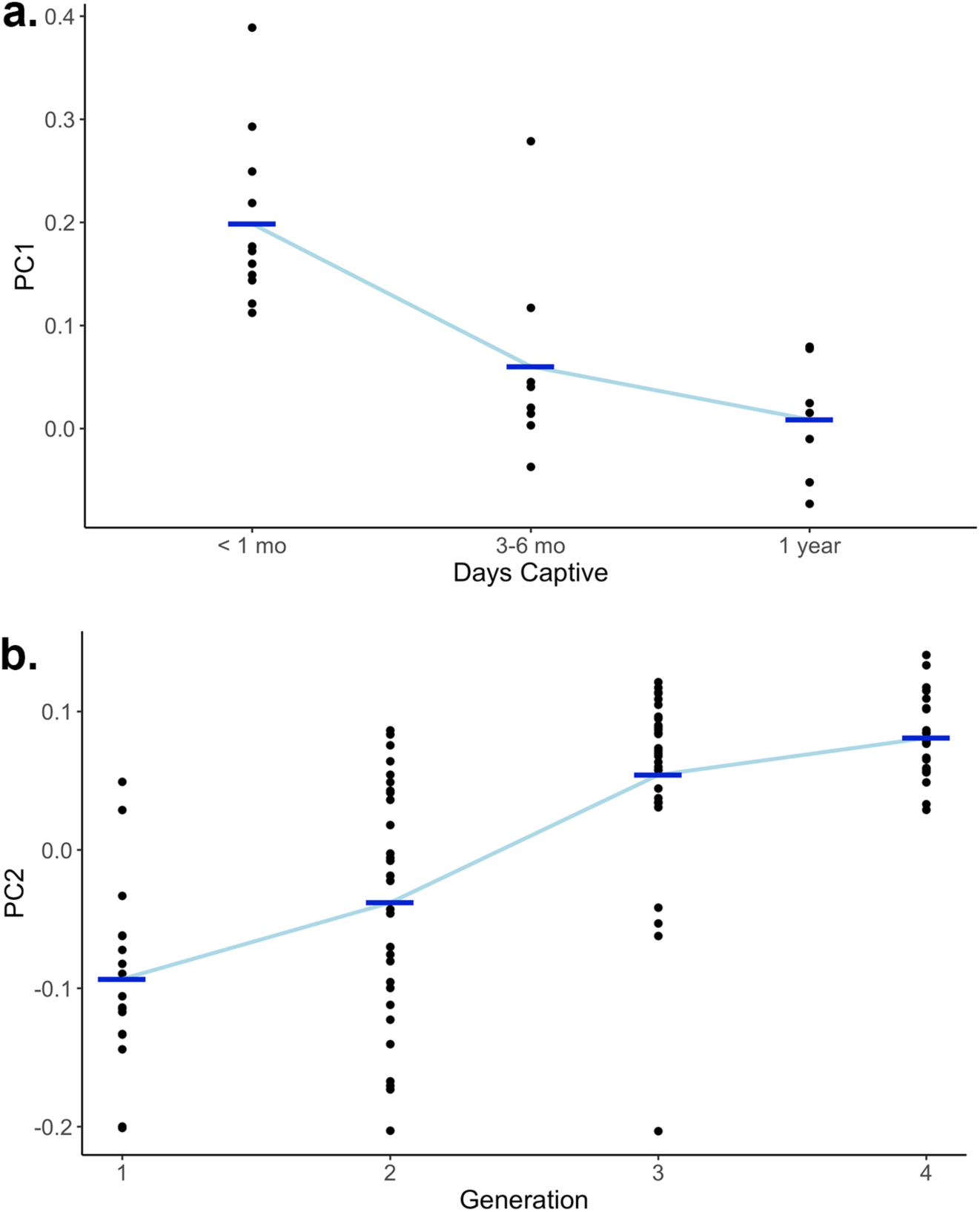
Different principal component axes (extracted from the ordination shown in Figure 1c) correspond to different time scales of change. Change along PC1 (a) is primarily driven by the number of days captive for F_0_, while change along PC2 (b) is primarily related to generation in captivity (F_1-4_).

**Table S1.** A randomly trimmed dataset provides a balance of sample observations across generations (See attached data file).

## Author contributions (CrediT taxonomy)

Conceptualization, C.L.W., S.N.D.K., and D.M.S.; Methodology, C.L.W., C.E.W., and D.M.S.; Software; C.L.W. and C.E.W.; Formal Analysis, C.L.W. and C.E.W.; Investigation, C.L.W., S.N.D.K., and D.M.S.; Resources, C.L.W. and D.M.S.; Data Curation, C.L.W., C.E.W., S.N.D.K., and D.M.S.; Writing-original draft, C.L.W. and C.E.W.; Writing-review & editing, C.L.W., C.E.W., S.N.D.K., and D.M.S.; Visualization, C.L.W. and C.E.W.; Supervision, C.L.W. and D.M.S.; Project administration, C.L.W. and D.M.S.; Funding acquisition, D.M.S.

## Acknowledgements

All research was permitted by the U.S. Fish and Wildlife Service, the California Department of Fish and Wildlife, and the San Diego Zoo Wildlife Alliance (SDZWA) Institutional Animal Care and Use Committee. This work was funded by California Traditional Section 6 grants to D.M. Shier and R.R. Swaisgood, the U.S. Fish and Wildlife Service, and a cooperative agreement N62473-20-2-0016 from the U.S Navy.

We would like to thank SDZWA staff and volunteers for their assistance in sample collection (M. Swartz, R. Chock, T. Wang, A. Kozuch, M. La Cava, S. Leivers, E. Drum, C. Dyslin), organization (Erin Drum, Eva Newby), and analysis (Marissa Taylor, Rachel Felton, Cynthia Steiner), and Aspen Reese and Jessica Diaz for their contributions to sample selection in the study design. We would also like to thank Alison Greggor for her assistance in compiling reproductive fitness measurements and Louis Gunning for assistance in processing pedigree data. We are extremely grateful to the entire Pacific pocket mouse team, past and present (M. Swartz, R. Chock, A. Kozuch, M. La Cava, J. Chang, A. Heath, T. Wang, S. Leivers, E. Drum, C. Dyslin, R. Gosselin, A. Greggor, A. Flanders, A. Harris and M. Moore), Aryn Wilder, and Christopher Tubbs for their insightful discussions and support.

PMx software (Ballou et. al. 2022) was provided under a Creative Commons Attribution-No Derivatives International License, courtesy of the Species Conservation Toolkit Initiative (https://scti.tools).

## Notes

### Competing Interest Statement

The authors have declared no competing interest.

